# Stereotypical modulations in dynamic functional connectivity explained by changes in BOLD variance

**DOI:** 10.1101/126524

**Authors:** Katharina Glomb, Adrián Ponce-Alvarez, Matthieu Gilson, Petra Ritter, Gustavo Deco

## Abstract

Spontaneous activity measured in human subject under the absence of any task exhibits complex patterns of correlation that largely correspond to large-scale functional topographies obtained with a wide variety of cognitive and perceptual tasks. These “resting state networks” (RSNs) fluctuate over time, forming and dissolving on the scale of seconds to minutes. While these fluctuations, most prominently those of the default mode network, have been linked to cognitive function, it remains unclear whether they result from random noise or whether they index a non-stationary process which could be described as state switching.

In this study, we use a sliding windows-approach to relate temporal dynamics of RSNs to global modulations in correlation and BOLD variance. We compare empirical data, phase-randomized surrogate data, and data simulated with a stationary model. We find that RSN time courses exhibit a large amount of coactivation in all three cases, and that the modulations in their activity are closely linked to global dynamics of the underlying BOLD signal.

We find that many properties of the observed fluctuations in FC and BOLD, including their ranges and their correlations amongst each other, are explained by fluctuations around the average FC structure. However, we also encounter interesting characteristics that are not explained in this way. In particular, we find that the brain spends more time in the troughs of modulations than can be expected from stationary dynamics.

## 1 Introduction

In their seminal paper, Biswal et al. (1995) report that correlation patterns in motor cortex during rest are remarkably similar to those found during a finger tapping task. Following this discovery, a number of large-scale functional networks was unveiled using the experimental paradigm of “resting state” (RS) and the concept of functional connectivity (FC) (Lowe et al., 1998; Cordes et al., 2000; Kiviniemi et al., 2003; Fox et al., 2005; Beckmann et al., 2005; De Luca et al., 2006; Damoiseaux et al., 2006; Mantini et al., 2007; Smith et al., 2009; Yeo et al., 2011). Investigating these functional networks, termed “resting state networks” (RSNs), enables us to probe integration (within networks) and segregation (between networks) in the brain.

More recent research has shown that the FC between brain regions forming the basis of RSNs are not constant over time. Dynamic functional connectivity (dFC) is defined in contrast to average or “static” functional connectivity (avFC), and dFC often appear to greatly deviate from avFC. Changes in the global FC structure apparent in dFC have been taken to be signs of time-dependent integration and segregation between functional networks (see Hutchison et al. (2013); Calhoun et al. (2014); Preti et al. (2016) for reviews). A growing number of studies are proposing and studying new and innovative approaches to characterizing dFC (Chang and Glover, 2010; Britz et al., 2010; Kiviniemi et al., 2011; Hutchison et al., 2012; Allen et al., 2012; Leonardi and Van de Ville, 2013a; Liu et al., 2013; Zalesky et al., 2014; Damaraju et al., 2014; Karahanoğlu and Van De Ville, 2015; Betzel et al., 2016a). Many of these studies have focused on characterizing recurring “states” of brain connectivity. This way, it was established that changes occur on the level of large-scale configurations of inter-network relationships, where some regions belonging to attention networks and the DMN act as “hubs” not only in terms of their static connectedness, but also in terms of the variability of their connections. This argues against the rigid assignment of brain regions to a functional network, and stresses that at certain points in time, the FC patterns can diverge greatly from the average.

Since the beginning of RS research, it was asked whether the observed fluctuations are “meaningful” or not. This is tightly related to the question of what the mechanism behind the fluctuations is. It has already been recognized that several different sources likely contribute to the variability observed in dFC (Messé et al., 2014; Barttfeld et al., 2015): First, the observed fluctuations can be shown to be relevant to behavior (Fox et al., 2005; Bassett et al., 2011; Kucyi and Davis, 2014; Yang et al., 2014; Davison et al., 2015) which has recently lead to the idea of “controllability”, i.e. that the large-scale architecture of the human brain is organized in such a way that frequently visited states are reachable with minimal effort and maximal energy efficiency (Gu et al., 2015). Thus, observed dynamics are partially linked to ongoing cognition and are implied to arise from temporal dynamics on a neural level (Shmuel and Leopold, 2008; He and Raichle, 2009; Schšlvinck et al., 2010; Thompson et al., 2013; Kopell et al., 2014). Second, a certain part of the fluctuations is related to changes in brain state, like arousal (Chang et al., 2013), sleep state (Larson-Prior et al., 2009; Horovitz et al., 2008), or aneasthesia (Vincent et al., 2007; Bettinardi et al., 2015; Hutchison et al., 2013). Third, recent work shows a link between ongoing fluctuations in the brain and rhythms generated in internal organs like the heart and stomach, suggesting that self-referential behavior related to the default mode network includes the body as well as the mind (Babo-Rebelo et al., 2016; Richter et al., 2016; Babo-Rebelo et al., 2016). Fourth, a part of the fluctuations arises from random noise reverberating through a structured network (Deco and Jirsa, 2012; Messé et al., 2014; Ponce-Alvarez et al., 2015; Glomb et al., 2017). Nonetheless, this last source of variability could be functionally relevant to the maintenance of synaptic connections. Furthermore, these reverberations constitue a large part of the broadband-signal (He, 2014; Chaudhuri et al., 2016), and only recently are receiving increased attention.

Evidently, the mechanisms behind dFC are manifold and require careful interpretation of data as well as integration of results from many lines of inquiry and different methods (Ritter et al., 2013). In the present study, we relate our results to the mathematical concept of stationarity. Observed data are generated by a random process, *x*_*t*_ = *y*, i.e. the data *y* are realizations of the random process {*X*}, according to some probability distribution which assigns a value to the random variable *x*_*t*_ at each point in time. For a weakly stationary process, the random variables exhibit the following properties:

1. constant mean: 𝔼(*x*(*t*)) = *μ*
2. constant variance: var(*x*(*t*)) = σ^2^
3. covariance is a function of time shift, not of time itself: cov(*x*_*t*_, *x*_*t*+*τ*_) = *f*(*τ*)

In other words, the first two moments of the probability distribution of *x*_*t*_ are required to remain unchanged over time. This means that correlations found in shorter time windows are bounded by long-term correlations. Of course, these distributions are not directly observable and must be estimated from the data. Thus, two tools whose importance cannot be stressed enough in this context are the usage of null data in the form of surrogates on the one hand (Handwerker et al., 2012; Bassett et al., 2013; Lindquist et al., 2014; Hindriks et al., 2015; Betzel et al., 2016a; Laumann et al., 2016), and modelling approaches on the other (Messé et al., 2014; Hansen et al., 2014; Glomb et al., 2017). The former allows us to construct stationary time series with the means, sigmas, and covariances of the data. The latter provides a situation where the underlying random process is known. Comparison of simulated with empirical data provides the interpretation of the observed variability as a “dynamic repertoire” of the brain (Deco et al., 2013). Analyzing this repertoire is crucial for understanding the brain in terms of general principles of information processing, pattern formation, integration across temporal and spatial scales, etc.

The temporal dynamics of modulations present in time courses of spontaneous brain activity measured with fMRI BOLD are the focus of the present study. We will first characterize these dynamics on the level of RSNs and the underlying BOLD signal, and then investigate in how far our findings are explained by stationary dynamics. To this end, we will use both stationary surrogate data and a stationary model.

## 2 Methods

### 2.1 Data

We use data previously published in Schirner et al. (2015), recorded at the Charité University Medicine in Berlin, Germany. Three datasets are recorded from each of 49 healthy subject aged 18 to 80 years (mean 42 years, 30 females, 19 males): 1) 22-minute (*T* = 661 frames, TR=2s) RS scans (eyes closed), 2) diffusion-weighted (dw) MRI for extracting structural connectivity (SC), 3) anatomical scans (T1-weighted).

Preprocessing of the BOLD data (dataset 1) includes high-pass filtering (Gaussian high-pass temporal filtering, FWHM 100s, as available in FSL), motion correction, linear registration to MNI space, and registration to T1-weighed images obtained from dataset 3. Dataset 3 is segmented using FREESURFER and parcellation is performed according to the Desikan-Killiany atlas (Desikan et al., 2006). Parcellation results are inverted from anatomical to functional space (dataset 1) in order to obtain mean time courses across all voxels of each region. Discarding the Corpus Callosum on both sides, we are left with *N* = 66 cortical areas.

DwMRI (dataset 2) was performed with 64 directions. For each voxel, the fiber orientation distribution function was computed, which was then used by the probabilistic fiber tracking algorithm implemented in MRTrix (Tournier et al., 2004, 2007) to initiate tracts in the grey matter-white matter interface obtained as a 3D-volume (FREESURFER) and converted to MNI space (FSL, Matlab) from dataset 2. Tracts are prolonged until they reach a termination region. The fibers themselves are not visible in the dwMRI output, as their thickness and their histological details are below imaging resolution. Hence, neither their thickness, nor their degree of myelination, nor any synaptic connections - all important factors if one wishes to compute the SC - are directly available. Furthermore, the number of tracts found between two locations depends on their distance and on crossing fibers, branching, etc., meaning that it does not necessarily reflect the strength of the pathway. In order to limit the impact of these confounds, connection strengths were assumed to be bounded by the area available at the grey matter-white matter interface, as well as the potential connection strength it could provide which was assumed to be the same (relatively) for all tracts terminating in that voxel. (see Schirner et al. (2015) for details). The number of streamlines detected per connection was then used as input to an algorithm which derives effective connectivity from structural connectivity using a noise diffusion model (see section 2.6).

### 2.2 Motion artifact removal

There is clear evidence showing that motion artifacts can contaminate measures of dynamic FC (Van Dijk et al., 2012; Power et al., 2012; Satterthwaite et al., 2013) even after motion correction during initial preprocessing. We follow the approach described in Power et al. (2012), shown to be quite effective in Satterthwaite et al. (2013). A combination of framewise displacement (FD) and a measure for global change in the BOLD signal, termed DVARS, is used. FD is based on the six rigid body movement parameters, three for translation (*d*_*x*_, *d*_*y*_, *d*_*z*_) and three for rotation (*α*, *β*, *γ*, converted from radians into mm using a radius of 50mm). It is defined as

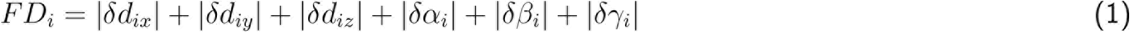

*δ* stands for the difference between frames *i* – 1 and *i*. DVARS is purely derived from the BOLD signal and is defined as

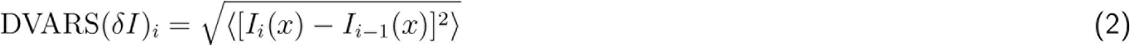

*x* is the vector of BOLD intensities at each ROI and angular brackets indicate averaging over ROIs.

We have to adapt DVARS to our purposes because its scale depends on the normalization of the BOLD signal. In Nichols (2017), the author derives a simple improvement on the DVARS measure which allows an expression in units of expected DVARS through normalization. Then, DVARS^⋆^ is defined as

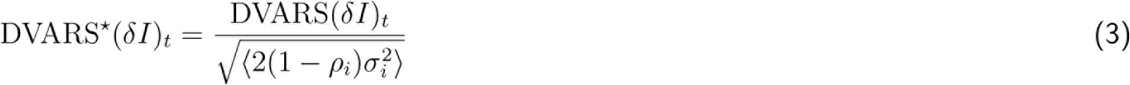

*ρ*_*i*_ is the the average autocorrelation between successive frames (*t*, *t* – 1) at ROI *i*, and *σ_i_* is the standard deviation of ROI *i*’s time course. The angular brackets denote averaging over ROIs. Since O is possibly itself contaminated by outliers caused by motion, it is approximated by the interquartile range. As a result, DVARS^⋆^ is in units of expected DVARS and is easily interpreted.

These two measures are complementary and frames which coincide with movement and exhibit unusually high BOLD changes are removed. However, for both measures it is necessary to choose a threshold. It is our goal to remove the relationship between motion parameters and dFC, and since the methods for motion correction used here were conceived for the voxel-level, we choose the thresholds according to a data-driven measure: we control the correlation between FD and instantaneous BOLD variance (see section 2.4). We use this measure instead of the other two as it is closely related to DVARS and therefore most likely to be influenced by it. The maximum correlation between between two subjects’ FD time courses - i.e. the random correlation - is 0.29. However, applying a sliding window is equivalent to low-pass filtering and as a result, this correlation increases up to almost 0.6, meaning that spurious correlations are introduced. Therefore, any motion artifacts will be smeared out in time and as a result, it is quite probable that unrelated motion artifacts appear to be aligned with an RSN or dFC measure windowed time course. Unfortunately, it is impossible to tell apart such cases from instances where a motion artifact is truly aligned with our time courses of interest. For DAVRS^⋆^, using 1 is an obvious choice, but since *ρ* and *σ* are still estimated from the presumably contaminated data, we considered thresholds ranging from 0.7 to 1.0. For FD, the value recommended in Power et al. (2012) is 0.5mm, but we found this to be quite high and tried values ranging from 0.1 to 0.5. Specifically, our procedure looks as follows:

1. compute FD from the motion parameters, DVARS^⋆^ from the BOLD signal
2. identify frames for which both FD and DVARS^⋆^ exceed their thresholds, hinting motion artifacts
3. replace these frames with NaNs (“scrub”)
4. using a given window width, compute instantaneous BOLD variance from the scrubbed data, ignoring the removed frames, as well as windowed FD by averaging FD inside each window
5. compute the correlation between instantaneous BOLD variance and windowed FD

Thus, for each parameter combination (combinations of thresholds, different window lengths, with and without global signal regression, see section 2.7) we find the subjects which have a correlation between their motion parameters and their dFC that exceeds the threshold, hinting at the possibility that motion artifacts remain after scrubbing. We then pick for each subject the highest (i.e. most generous) thresholds that lead to removal of motion artifacts, i.e. a correlation between FD and instantaneous BOLD variance of < 0.29. If for any subject there is no combination of threshold values which will remove the artifacts while keeping at least 70% of the frames for both window lengths (Satterthwaite et al., 2013), with and without GSR, we remove this subject. This is the case for 21 subjects, leaving us with 28 datasets. It should be stressed that this does not mean that the data are of poor quality. It is rather a strong reminder that the sliding window technique has its limitations. Not all excluded subjects are actually contaminated by motion artifacts, but we only keep the ones for which we can be certain that they are not. There is no correlation between the residual correlation and the number of removed frames.

Table S1 (supplementary methods section S1.1) shows the remaining correlations and the percentage of removed frames for each subject included in the analysis. For surrogates, we simply remove, after phase shuffling (see section 2.5) the same frames as in the original data. For every processing step downstream from scrubbing, the NaNs are simply ignored. While this obviously introduces errors in the windows that contain many NaNs, the same procedure is applied to surrogates and thus, if there is any systematic bias, it would be present also there.

### 2.3 RSN time courses

We use tensor decomposition, a dimensionality reduction approach which has previously been shown to be useful in neuroscience (Beckmann and Smith, 2005; Cichocki, 2013; Leonardi and Van de Ville, 2013a; Leonardi and Van De Ville, 2013b; Ponce-Alvarez et al., 2015) and which we apply to community detection (Gauvin et al., 2014); in our case, the communities we wish to detect are recurring FC patterns. For each subject (or simulation), we compute pair-wise FC inside of sliding windows (figure 1A), resulting in *W* time windows. This results in an *N* × *N* dynamic FC matrix, *dFC* (*w*) with *w* = 1,2,…, *W*, which is symmetrical. We use a non-negative measure of dFC, i.e. mutual information as described in Kraskov et al. (2004) for estimating MI between time series *X* and *Y*, which are *M* BOLD measurements for two ROIs inside a given window:

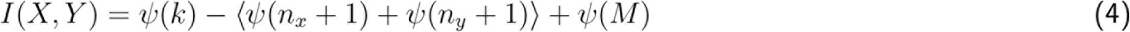

*ψ*(·) denotes the digamma function 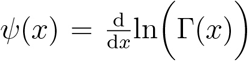. *k* =1 and denotes the usage of the nearest neighbor. This measure is based on counting the numbers *n*_*x*_ and *n*_*y*_ of points whose distance to a given point, projected on the x- and y-axis, respectively, is shorter than that to its nearest neighbor. The shortest distance is computed using the minimum norm

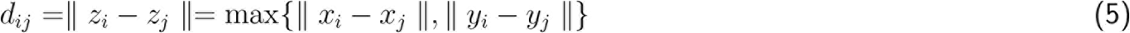

This is done for each pair of ROIs in each time window, resulting in *W* dFC-matrices. These *W* dFC-matrices are then stacked along the temporal dimension into 3-way data structures of dimensions *N* × *N* × *W*, called tensors (figure 1B) and denoted by <u>**Y**</u>. The reason for choosing to use MI instead of correlation is that we use non-negative canonical polyadic decomposition (NCP) to decompose our tensors (Kim and Park, 2012; Bader et al., 2015; Acar et al., 2011). NCP only accepts nonnegative values, and using MI avoids the difficulties that arise from interpreting negative correlations. Furthermore, we showed previously (Glomb et al., 2017) that MI leads to better clustering performance downstream in the processing pipeline (see below) than either correlation itself or absolute values of correlations.

It has been a long-standing problem in the field of dFC how to identify significant correlation values when dealing with a multitude of time windows, subjects, and pairs (Zalesky and Breaks-pear, 2015; Hindriks et al., 2015). Here, we assess the quality of features resulting from tensor decomposition to set the parameters of our decomposition procedure (see below). Instead of identifying significant dFC values, we threshold and binarize the tensors using different percentiles, i.e. *θ* = [0,75,80,90,91,…99]. For example, *θ* = 99 means that the 99% lowest dFC values inside a tensor are set to 0 and the remaining 1% to 1. At *θ* = 0, we keep the continuous values (no binarization).

**Figure 1:**
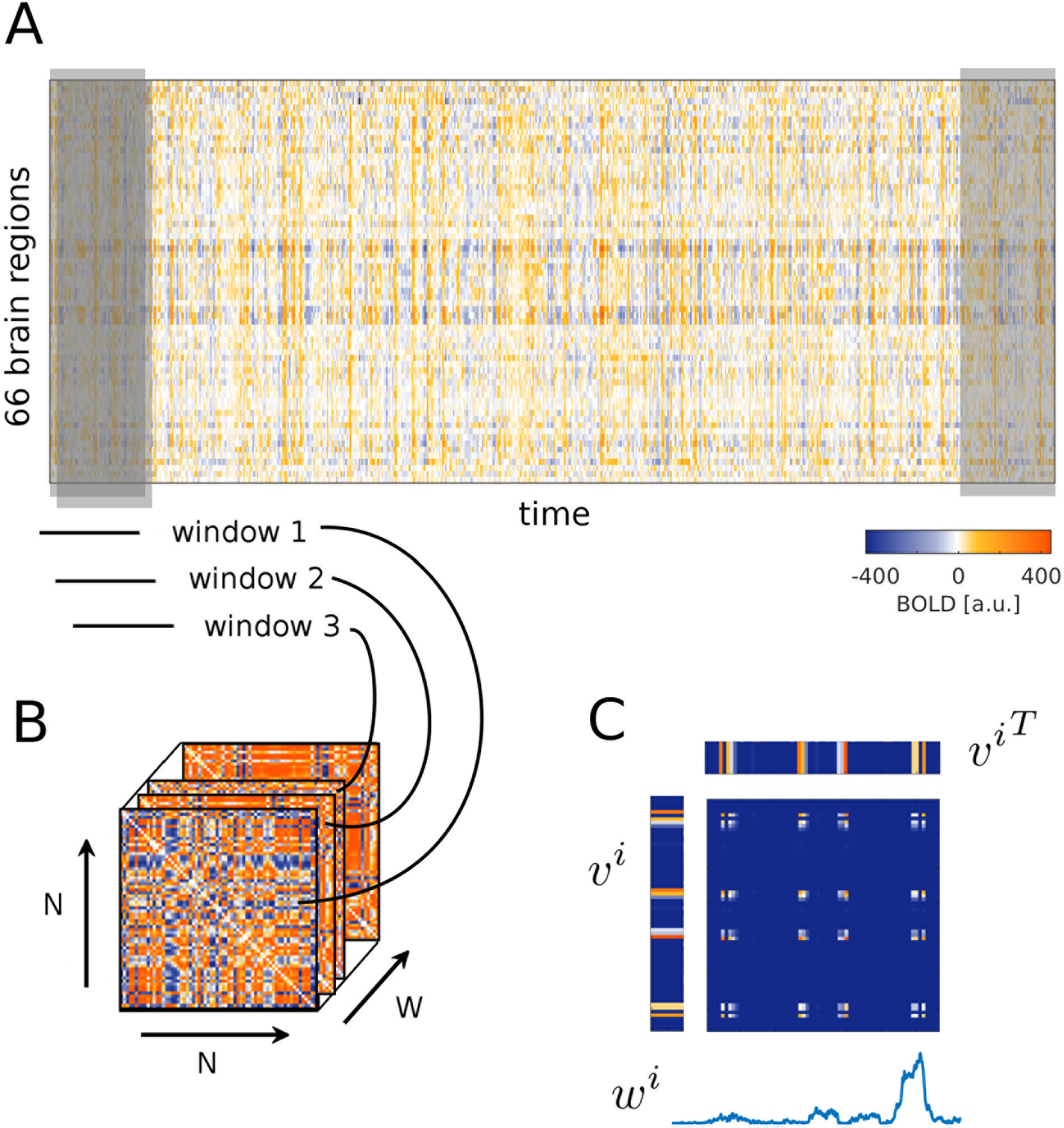
Illustration of sliding windows dynamic FC and tensor decomposition. **A** Example time courses for one subject. The slliding window are illustrated in grey: it is moved alond the entire time course, resulting in *W* windows, each of which covers a portion of the BOLD time courses. **B** The data points that fall into any given window are used to compute a dynamic FC matrix. DFC matrices from all *W* windows are stacked to form the tensor. **C** The tensor is decomposed into a given number *F* of sets of three vectors. Each set i contains two spatial features *v*^*i*^, i.e., the communities; they are identical due to the symmetry of the dFC-matrices. The third vector *w*^*i*^ is the time course of weights associated with this community. Taking the outer product 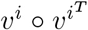 results again in a matrix. Any given dFC matrix is approximated by adding all *F* matrices given in this way for the tensor, weighed by the value *w*^*i*^(*t*).

Another parameter is the number *F* = [3,4, …9] of features that we extract from each tensor. Each feature i is a rank-1-tensor, i.e. it can be represented as the outer product, denoted as ○, between three vectors, each vector corresponding to one dimension of the tensor (figure 1C):

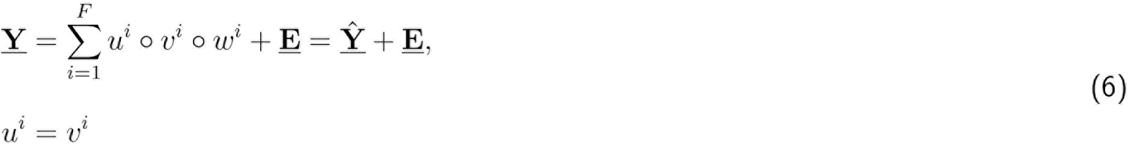

Since the dFC matrices are symmetric, the first two vectors (of length *N*) are identical (*u*^*i*^; = *v*^*i*^) and are interpreted as containing membership weights. These weights describe how strongly each of the 66 brain regions is a member of community *i*. The third vector *w*^*i*^ (of length *W*) is the time course which describes how strongly this community as a whole contributes to the dFC in each window. Of course, the decomposition which we obtain is only an approximation, leaving the variability contained in the error <u>**E**</u> unexplained. Decomposition fits are shown in SI methods, section S1.2, figures S1-S4.

The vectors containing the membership weights, or “spatial features”, are pooled across all subjects and clustered five times using K-means clustering with *K* = [3, 4, …10]. The reasoning behind this is that if our decomposition works and we indeed find RSNs, they should occur in many (if not all) subjects. This is a way to validate our results in a data-driven manner without relying too much on templates obtained from previous publications with different methods, which could bias our results and remove potential benefits of our approach. The cluster centers are then used as templates which can be compared to spatial features obtained from single subjects, surrogate data and simulated data. We assess the goodness of clustering via the silhouette value (de Amorim and Hennig, 2015), comparing to phase-randomized surrogate data (section 2.5) which do not contain a cluster structure as their long-term correlation structure has been destroyed. We choose the values of *θ*, *F* and *K* such that the difference between the sihouette values for surrogates and real data is maximal. Because of our coarse spatial parcellation, we do not require each network to occur in each subject. At the same time, it does not make sense to search for more RSNs per subject than there are clusters. As a result, for each value of *F*, we only use *K* ≥ *F*. We confirm that the results are robust by repeating the decompositions and subsequent clustering several times. The optimal parameter settings are remarkably reproducible. All silhouette values can be found in SI methods section S1.3 and figures S5-S8.

Thus, for empirical data, the RSN identity is its cluster membership. For surrogate and simulated data, we use confusion matrices to compute how similar extracted features are to the templates (comparing each community vector to each template). First, we quantize extracted features and templates on three levels and count how many times the levels match. This results in one confusion matrix per template-feature pair. Then, Cohen’s *κ* is computed as a summary measure of each confusion matrix:

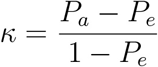

*P*_*a*_ is the overlap, *P*_*e*_ is the expected overlap based on the distribution of the levels, similarly to Hamming distances which can be thought of as using only two quantization levels. Each feature is assigned to a template according to its maximum *κ*-value. A perfect overlap woult result in *κ* =1. For simulations, an average *κ* is computed for different values of the model parameter *G* (global coupling, see section 2.6) and the *G* with the maximum *κ* is chosen. Details can be found in SI methods section S1.4, figure S9 and table S2.

### 2.4 Measures of RS dynamics

We use three measures to quantify global dynamics of RS. Two measures are related to changes in functional connectivity (FC), and one to changes in the BOLD signal itself. For each of them, one time course is created for each subject, using the same sliding windows as for the tensors in order to be able to relate the resulting time courses to the RSN time courses.

The FC-related measures rely on pairwise correlations, i.e. as for the tensors (section 2.3), we compute dFC-matrices for each window, containing pair-wise FC values, but this time, using correlation instead of mutual information. This may seem like a contradiction, but in fact, results are more reliable if they do not depend on the exact FC measure. Again, the dFC matrices are symmetrical, and their diagonals are filled with ones. The *instantaneous average correlation* is just the average over all unique pairs, i.e. using the upper or lower triangle of the *dFC*(*w*) matrix, excluding the diagonal:

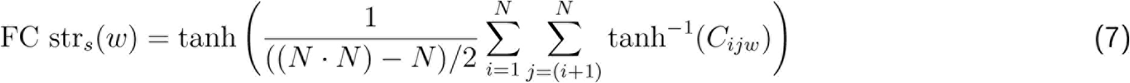

*C*_*ijw*_ is the entry in the *dFC*(*w*) matrix at pair *ij* and window *w*, *s* is the subject index, tanh^−1^ is the hypoerbolic arctangent, needed to z-transform the correlation values before averaging them, and likewise, tanh is the hyperbolic tangent, needed to back-transform the result into a correlation value. In other words, this is the average FC present in each slice of a tensor (without thresholding).

We evaluate how similar each *dFC*(*w*) matrix is to the average FC (avFC) matrix, i.e. the FC matrix obtained from correlating the full time courses of the ROIs. We call this measure *similarity to avFC*. This is done by simply correlating the upper or lower triangle of the matrices with each other after flattening them into a vector.

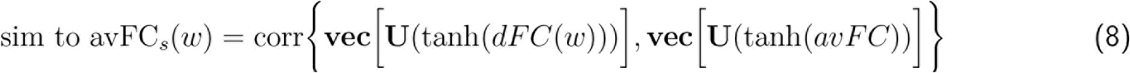

**vec**(·) stands for the vectorization of a matrix, **U** is the upper triangle of a matrix, and corr stands for Pearson correlation.

Additionally, we use a measure that tracks changes in the BOLD signal itself, namely of its variance. For this, a normalization step is necessary because the variability of variances across subjects does not carry any meaning, only the relative differences between ROIs within a subject do. Therefore, for each subject, we normalize such that the variance of its most variable ROI is 1. We then compute the variance of the signal x_*i*_ for each ROI *i* inside each window of length *w* ranging from *t* to *t* + *w* – 1 (in frames) and average over ROIs, resulting in a measure for the mean variance of the signal, which we refer to as instantaneous *BOLD* variance.

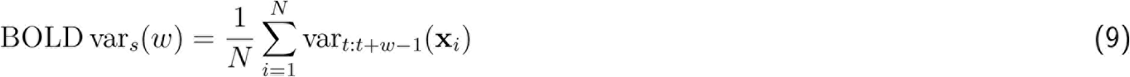

Note that this is not the variance of the mean signal.

### 2.5 Surrogates

Surrogate data are constructed by randomizing an aspect of the empirical data with the goal to produce data that preserve all characteristics except the one of interest. In this case, this characteristic is nonstationarity, corresponding to the null hypothesis that observed dynamics are a result of the long-term correlations, but that no additional structure is present. Correlation is closely related to covariance and thus to autocovariance, which are all second order statistics and required to be constant in time in a weakly stationary process. The representation of the autocovariance function in frequency space is the power spectral density. Thus, creating a situation in which this density is constant over time is equivalent to obtaining constant correlation. In order to do just this, we use the Fast Fourier Transform (FFT) to obtain a representation of the signal in frequency space and then add random phases to each bin.

In particular, we have *x*_1_, *x*_2_, …, *x*_*T*_ and *y*_1_, *y*_2_, …, *y*_*T*_, where x and y are two ROI’s signals in time whose long-term correlation we want to preserve. After applying the FFT and obtaining signals *X*_1_, *X*_2_, …, *X*_*N*_ and *Y*_1_, *Y*_2_, …, *Y*_*N*_ in frequency space, a random phase vector *φ*_*r*,1_, *φ*_*r*,2_, …, *φ*_*r*,N_, values of which are drawn from a uniform distribution in the interval [–*π*,*π*], is used to randomize phases in each frequency bin. The long-term correlations are preserved by adding the same phases to both signals before transforming back. For the surrogates that are used to determine the cluster performance, long-term correlations are not preserved and phases are added individually.

### 2.6 Model and simulations

We use a dynamic mean-field (DMF) model (Wong and Wang, 2006; Deco et al., 2014) where each of the 66 cortical regions are modelled by a node. Each node *i* contains one excitatory and one inhibitory pool which are described by their population firing rates *r*_*i*_, currents *I*_*i*_, and synaptic gating variables *S*_*i*_ according to the following coupled differential equations:

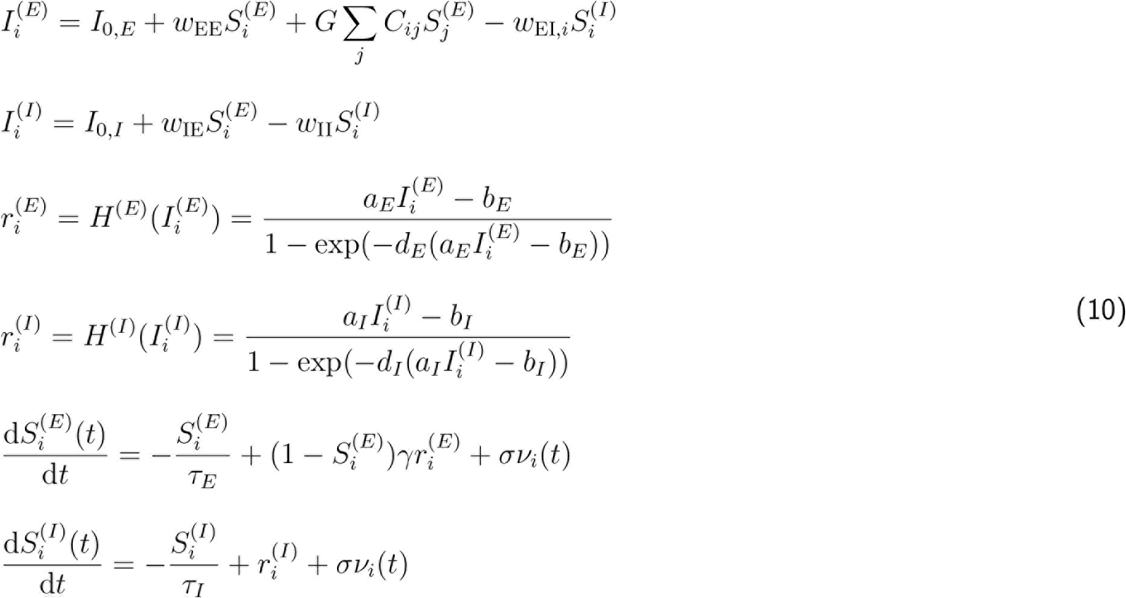

The model is illustrated in figure 2A. Locally, excitatory (indexed by *E*) and inhibitory (indexed by *I*) pools are connected to each other, using weights *w*_EE_, *w*_EI,*i*_, *w*_IE_, and *w*_II_. Most of these weights are constant (*w*_*EE*_ = 0.21, *w*_*II*_ = 1, *w*_*IE*_ = 0.15), except for the feedback inhibition *w*_EI,*i*_ which is adjusted prior to running simulations in order to keep mean firing rates in each population between 3 and 10 Hz. This is necessary because in this asynchronous regime, the model only has one attractor and is thus stationary (Deco and Jirsa, 2012). This model includes three types of synapses: 1) NMDA for local excitatory connections with *τ*_*E*_ = 100 ms, 2) GABA for local inhibitory connections with *τ*_*I*_ = 10 ms, and 3) AMPA for long-range connections between excitatory pools, governed by the connectivity matrix **C** with entries *C*_*ij*_ setting the weights between nodes *i* and *j*. The dynamics of these synapses are not modelled directly because they reach their steady state almost instantaneously (Deco et al., 2014). Additionally, each population receives a constant background current, *I*_0,*E*_ = 0.382 nA and *I*_0,*I*_ = 0.267 nA. *C*_*ij*_ are entries in the adjacency matrix governing the weights between the nodes. These weights are scaled by the factor *G*, which is a global coupling parameter, and the only free parameter of the model.

Here, C is the effective connectivity (EC) matrix, shown in figure 2B, derived by combining DTI-based structural connectivity (SC) with a noise diffusion (ND) model (Gilson et al., 2016). Most models that relate structure and function in RS fMRI use only the zero time shift-covariances to fit model parameters. As a result, directionality cannot be recovered because these covariance matrices are symmetric. Here, each connection weight of the SC (averaged over all subjects) is adjusted via a Lyapunov optimization procedure in order to fit the average “conventional” (zero time shift) as well as the time shifted covariance matrices, resulting in a weighted and directed graph. We use only the strongest anatomical connections identified by fiber tracking, obtaining a connectivity of 32%.

We previously showed (Glomb et al., 2017) that, when the nodes in the DMF model are connected according to this matrix, communities can be extracted from the resulting dFC time courses that approximate the RSNs found in empirical data. The EC matrix is asymmetric, contains homotopic connections, and exhibits a nearly uniform degree distribution which promotes stable simulations over a wide range of values of *G*. The results we show here are obtained by simulating with the value of *G* which leads to the maximum overlap with RSNs obtained from empirical data (see section 2.3).

**Figure 2:**
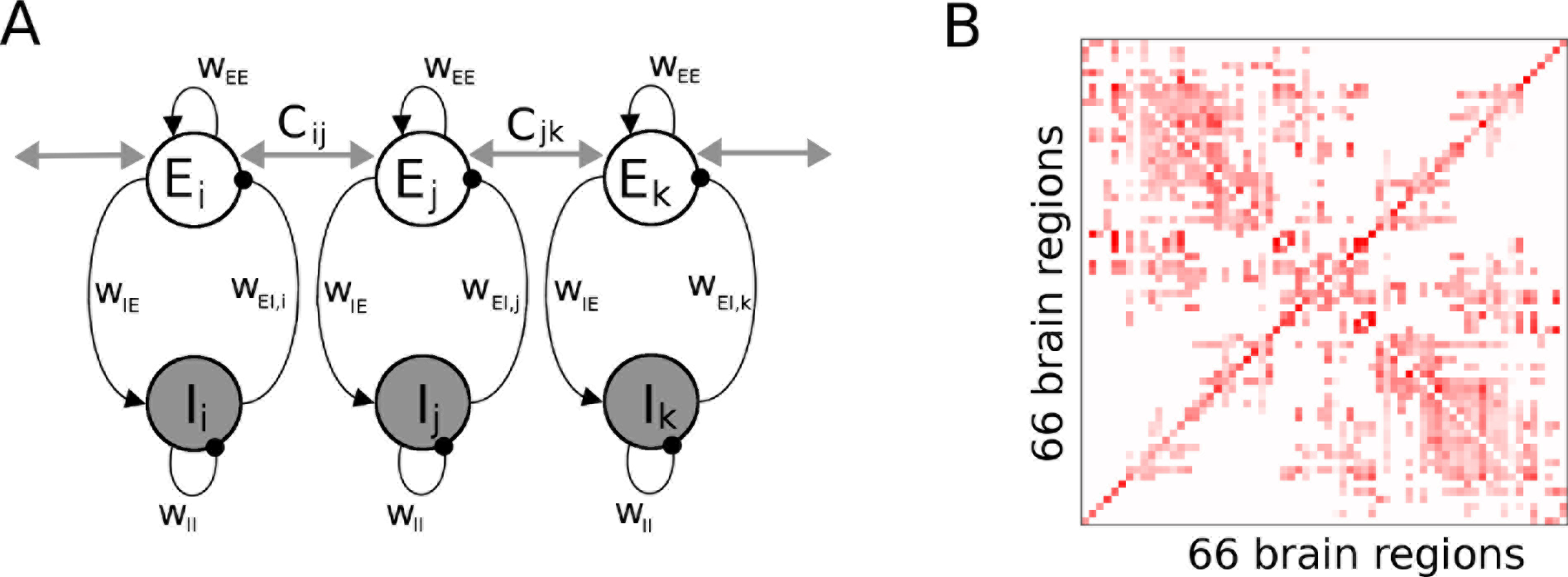
**A** Illustration of DMF model used to produce stationary simulated data. Each node contains an excitatory (E) and an inhibitory (I) pool which are locally connected by GABA (black lines, spheres) and NMDA (black lines with arrows) synapses and E pools have long range AMPA connections (gray lines) set by entries *C*_*ij*_ of the connectivity matrix. **B** Effective connectivity matrix. Note the asymmetry and the even distribution of the weight magnitudes. Weights are in arbitrary units.

### 2.7 Impact of window length and global signal regression

Throughout the result section, we use a window length of w=120s and we do not include global signal regression (GSR) in the preprocessing of the time courses. Although some studies apply this latter technique, it was shown by Saad et al. (2012) that it removes important information and distorts the correlation structure, including, but not limited to, the introduction of anticorrelations and changes in network properties (Murphy et al., 2009; Satterthwaite et al., 2013). However, in order to show that the effects demonstrated here are not attributable to a generalized global modulation in signal strength, we conducted the same analyses including GSR. Furthermore, while our window length is within the usual range (Leonardi and Van De Ville, 2015), it has been suggested that shorter window lenghts are also applicable (Zalesky and Breakspear, 2015). Likewise, it is certainly desirable to demonstrate that our results are not specific to our window length. Therefore, in each section, we mention these results. Details can be found in the supplementary information.

## 3 Results

### 3.1 Resting state networks coactivate strongly

One of the most prominent approaches to investigating time-dependent FC is to measure pairwise FC inside of overlapping sliding time windows, thus obtaining a sequence of dynamic FC matrices (dFCMs, figure 1A, B).

We were interested in shedding light on the basis of these fluctuations and their impact on RSN dynamics, given that it remains unclear whether RSNs are separate in time (Baker et al., 2014; Betzel et al., 2016a) or not (Mitra et al., 2014; Karahanoğlu and Van De Ville, 2015; Glomb et al., 2017). Figure 3 illustrates these two scenarios schematically.

**Figure 3:**
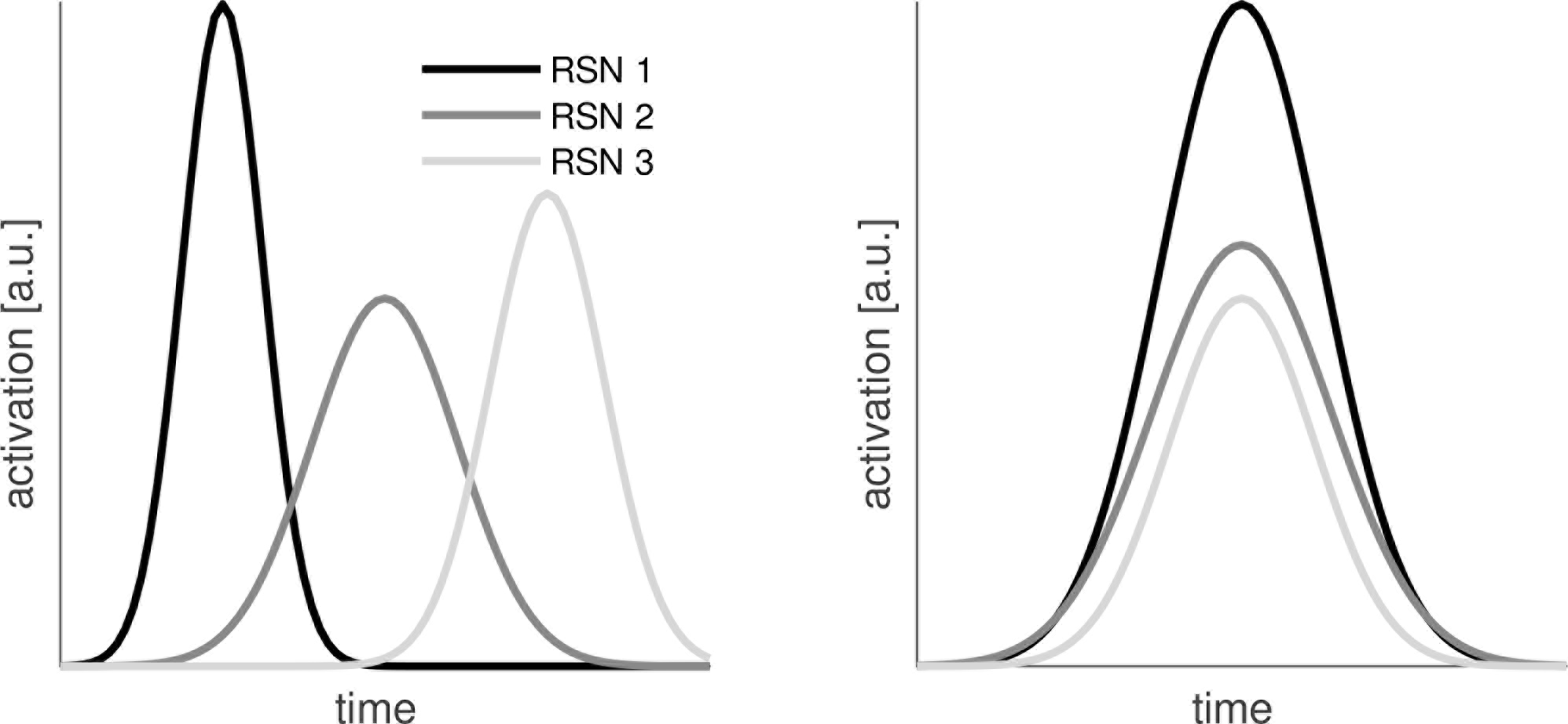
Illustration of two alternative scenarios for RSN activation time courses. (a) RSNs could activate in sequence. At any point in time, one RSN’s weight would be much higher than the other two. (b) RSNs could coactivate. At any point in time, combinations of RSNs would be present.

We use 22 minute long resting state (RS) scans from *S* = 28 healthy participants (Schirner et al., 2015), parcellated according to the Desikan-Killiany atlas (Desikan et al., 2006), obtaining *N* = 66 large-scale brain regions covering the entire cortex. We use nonnegative canonical polyadic decomposition (NCP), a highly interpretable version of tensor decomposition (Cichocki, 2013; Gauvin et al., 2014; Ponce-Alvarez et al., 2015), to obtain RSN time courses. This approach approximates a sequence of dynamic FC matrices (dFCMs) as a linear superposition of a small number of static FC patterns. The patterns’ weights and thus, the degree to which they contribute to dFC, are time-dependent. In contrast to the most frequently used approach, independent component analysis (ICA), these networks are allowed to overlap in both space and time. The extraction works as illustrated in figure 1: We compute pairwise dFC inside each time window, obtaining a series of dFCMs for each subject (figure 1A, B). Next, the dFCMs are thresholded and binarized with threshold *θ* to retain only the highest pairwise dFC-values. Each tensor is decomposed into *F* features (figure 1C), meaning that for each subject, we obtain *F* spatial and *F* associated temporal features, i.e., their activation time courses. The spatial features are vectors of length *N* = 66 (number of brain regions in our parcellation), the time courses are vectors of length *W* (number of time windows). After the decomposition, all subjects’ spatial features are pooled and K-means-clustering is applied. We use the cluster centers as “templates”; these templates resemble known RSNs (Glomb et al., 2017). As a result, we obtain a small number of RSNs that generalize across subjects (figure 4): somatomotor network, visual network, default mode network, control networks, and the dorsal visual stream, a component that is usually split into two lateralized ones, but is mixed at our level of resolution (Beckmann et al., 2005). For details, see section 2.3 and Glomb et al. (2017).

We repeat the analysis for a shorter sliding window (60s) and apply GSR before computing the tensors in order to establish the robustness of our procedure. Resulting RSNs are shown in figures S10-S12 (supplementary methods section S2.1), and are found to be very similar in each case. However, the clustering quality is better when not applying GSR, in line with Murphy et al. (2009); Saad et al. (2012). We conclude that GSR removes relevant information. The threshold selected by the procedures using GSR is the 99th percentile, effectively removing 99% of the data; when GSR is not used, this value is 98%.

**Figure 4:**
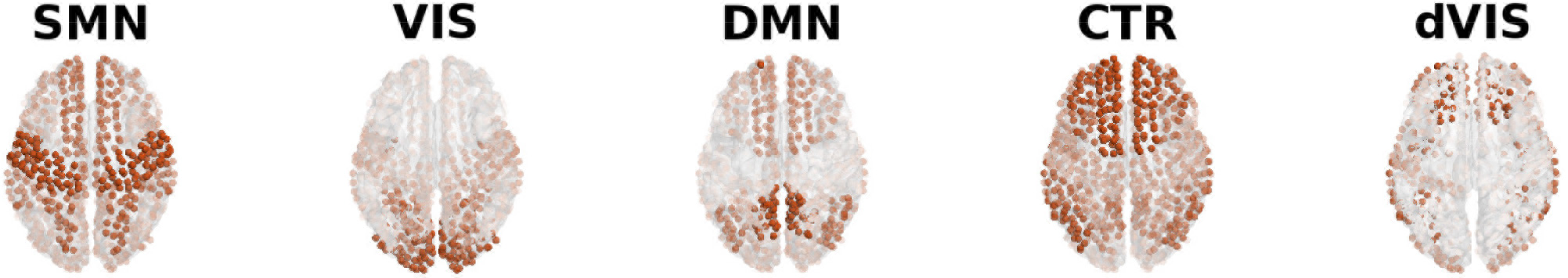
Community extraction from empirical data leads to recovery of RSNs. Each subject’s tensor is decomposed into three features, and the three community vectors are pooled across all subjects. Pooled community vectors are clustered. Using the cluster centers as templates and projecting them back to the cortex surface, we recover five known RSNs: SMN - somatomotor network, VIS - visual network, DMN - default mode network, CTR - control networks, dVIS - dorsal visual stream.

Note that for the tensors, we use mutual information (MI), computed following Kraskov et al. (2004), as pairwise FC measure (see section 2.3)s. We found in a previous study (Glomb et al., 2017) that MI is more reliable than correlation, at least at this spatial and temporal resolution, possibly due to its robustness towards outliers.

The associated time courses (figure 5) are interpreted as activation time courses. Our first observation is that the activations fluctuate considerably, and that these fluctuations seem to be strongly modulated on an infraslow time scale of 50 to 100 seconds. Note that the reason that all three time courses are zero at times is the thresholding applied to the tensors: there are time windows in which there is virtually no pair of brain regions that exhibits dFC above the threshold. This hints at dFC being clustered in time. Time courses for the other parameter settings can be seen in supplementary results section S2.2 and figures S13-S15.

The frequency of the fluctuations is not the focus of the present study since it is indeed hard to determine their properties when using the sliding window-approach. Instead, we investigated to what degree the RSNs overlap in time. The two scenarios depicted in figure 3 predict, respectively, that RSN time courses are dominated by transients, i.e. periods in which one RSN is active while the others are quiescent (left panel), or that instead, all/several RSNs’ time courses should be closely correlated (right panel). We consider the “contribution” of each RSN to the overall activation in each time window, which is just the ratio between its activation and the sum of all activations in this window. If this value is close to 1, this network dominates in the given time window. If the value is close to 1/*F*, i.e. 1/3 in this case, there is a large degree of co-activation; and if the value is very small, the network is not active at all.

Figure 6A shows the contribution values across windows for each RSN, pooled across subjects. The distributions show a prominent peak around 1/3, a value which corresponds to perfect coactivation in the case of having three RSNs per subject. There are also many zeroes, indicating windows in which the network was not active (windows in which all three were zero were excluded from the analysis since “contribution” cannot be defined in this case). The number of values falling in a higher range (right side of the plot) does not counter-balance this peak, suggesting that there is a bias towards RSNs being coactivated or not active at all and away from single RSNs being dominant. This prevalence of coactivation clearly contradicts the notion of independent networks. Inspecting histograms obtained for a shorter window and when using GSR (section S2.3, figures S16-S18) show that they exhibit the same basic properties.

**Figure 5:**
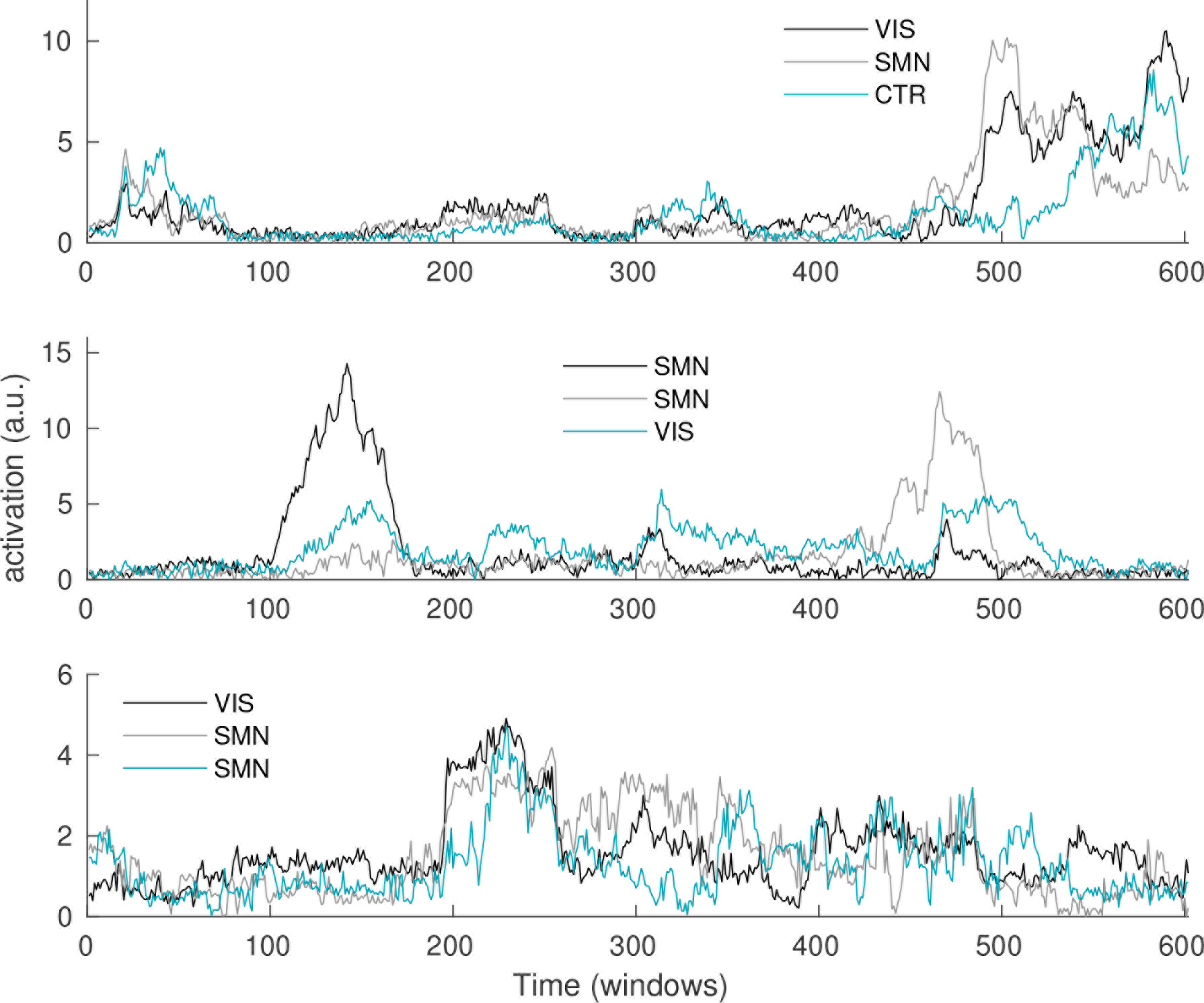
Example time courses of RSNs for three subjects. The time courses are obtained from the tensor decomposition and describe the degree to which each network contributes to the dFCM in each window and thus, how much it is activated.

**Figure 6:**
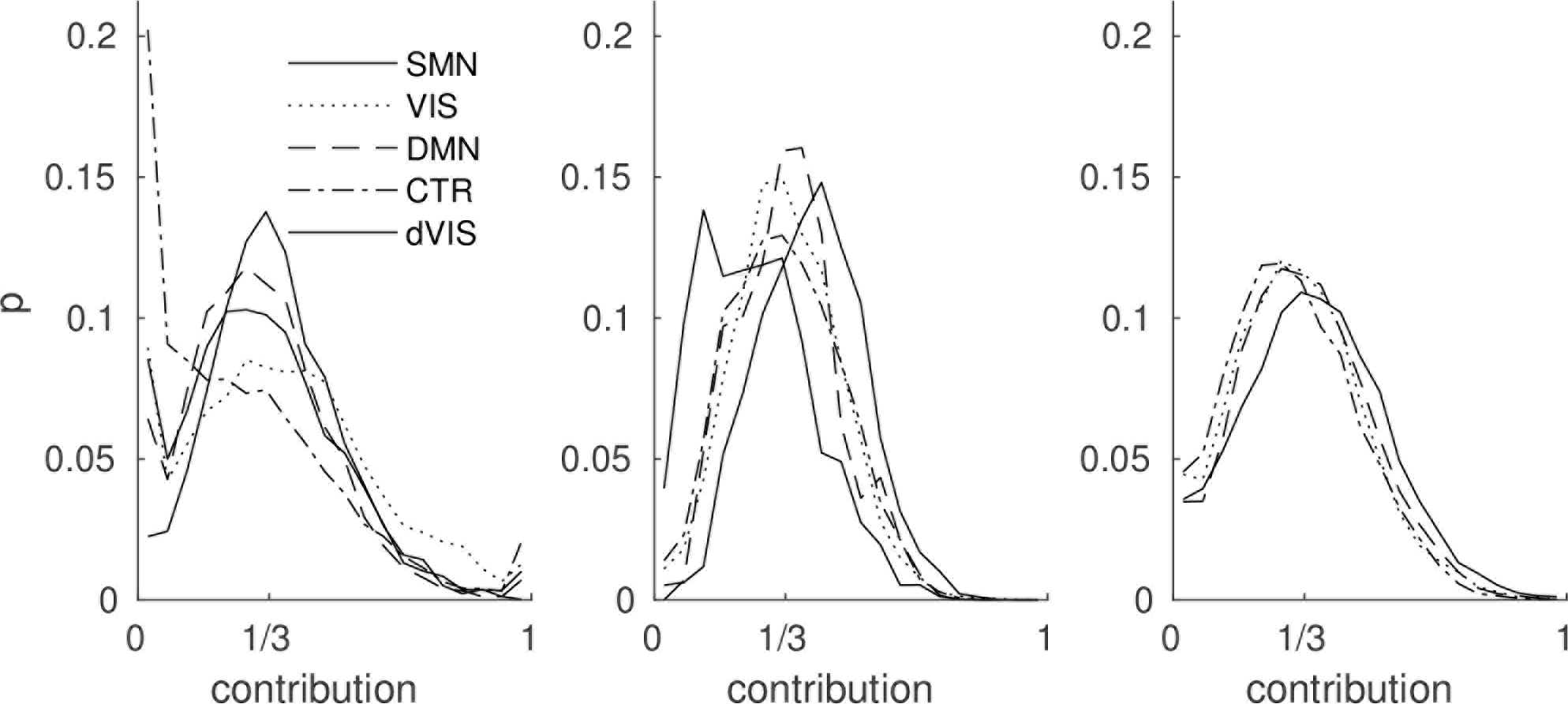
Contribution histogram for each RSN, pooled across subjects. Left panel: Empirical data, middle panel: surrogate data, right panel: simulated data. Since there are three features per subject, a value of 1/3 means perfect coactivation between the three RSNs. Higher values correspond to dominance of one RSN over the others. Values close to 0 indicate deactivation. The large peak around 1/3 clearly favors the coactivation scenario in figure 3. SMN - somatomotor network, VIS - visual network, DMN - default mode network, CTR - control networks, dVIS - dorsal visual stream

### 3.2 Modulations in RSN activation driven by underlying BOLD

In order to explain the observation that RSNs coactivate strongly, we take a step back and look at the underlying BOLD signal. Considering the example shown in figure 1A, one immediately notices “bands” or “events” as many, if not all, ROIs increase or decrease their BOLD activity simultaneously. It is fair to assume that these strong modulations will have an effect on the overall FC strength (Chawla et al., 1999; Daffertshofer and van Wijk, 2011) and eventually, on RSN dynamics themselves. We quantify this with two simple global measures. First, we take the variance of each ROI’s BOLD signal, averaged over all ROIs, inside each window. In time windows when all ROIs increase or decrease their activity together, this value will be high. If there are points in time where there are strong changes in different directions (i.e. some ROIs increase, others decrease in activity) this will likewise be captured by this measure. We hypothesize that big changes will have an impact, no matter their directionality. We term this the *instantaneous BOLD variance* (figure 7, orange). Second, we take the average over all pairwise correlations in each window (figure 7, blue) and term this *instantaneous average correlation* (see section 2.4).

It is immediately visible that these two measures are closely correlated. We compute the correlation coefficient on the group level by normalizing and pooling all subject’s time courses, obtaining a value of 0.68. On the single subject level, we obtain an average of 0.77 with standard error (SE) of 0.42. The reason that this average is higher than the value obtained from the group is probably that applying the sliding window introduces spurious correlations between subject time courses, leading also the dFC measures to be not independent between subjects. The window length does not have an impact on these numbers while GSR reduces them. Still, even when using both a smaller window and GSR, we still obtain a group correlation of 0.52 (0.62 with SE=0.32 on the single subject level), showing that our results are robust against window length and GSR. Results for all parameter setting can be found in section S2.4, table S3, and examples similar to figure 7 are shown in figures S19-S21. These considerably sized correlations suggest that there is an immediate need for methods that disentangle the effects of variations in signal strength on FC.

**Figure 7:**
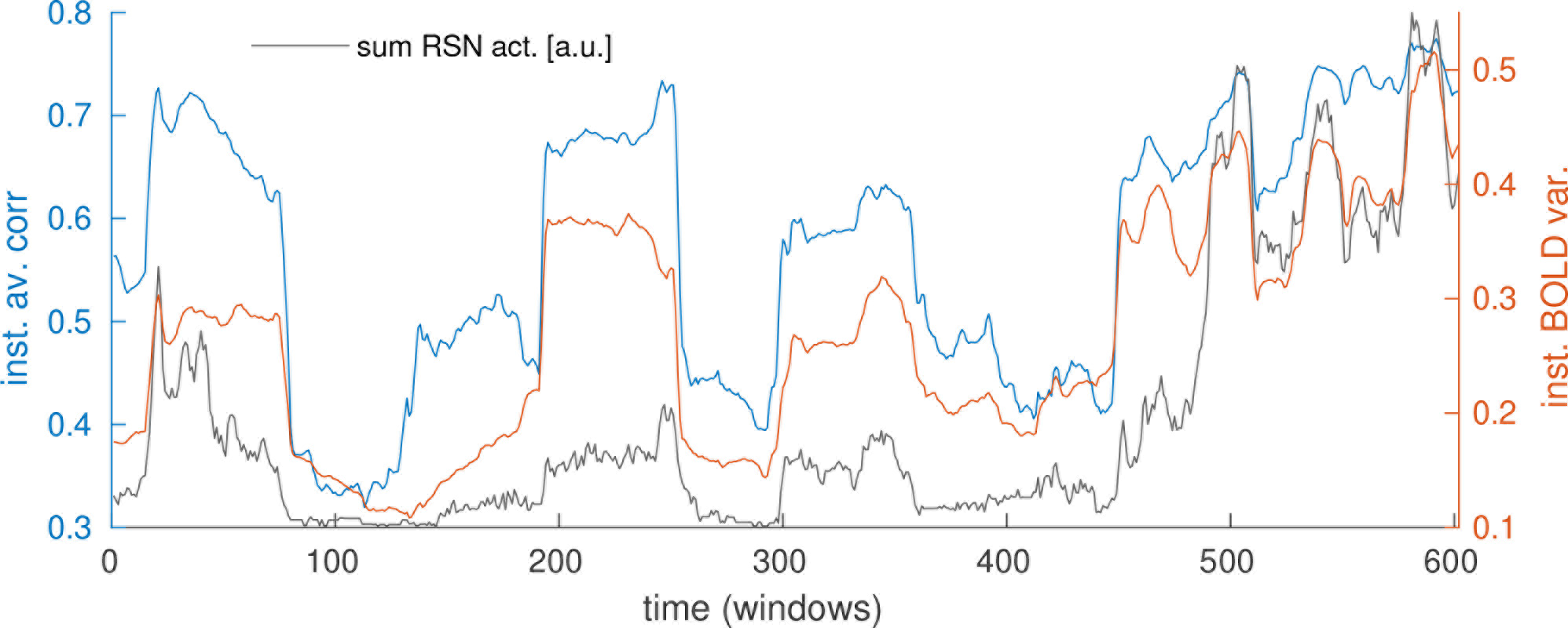
Global modulations in underlying BOLD signal (blue and orange curves) are tightly correlated with RSN dynamics (grey curve). Orange: *instantaneous BOLD variance*, blue: *instantaneous average correlation*, grey: sum over all RSN activations.

Taking this a step further, we also correlate the instantaneous BOLD variance with the previously extracted RSN time courses. We find a correlation of 0.74 on the group level, and 0.81 (SE=0.40) across all subjects, indicating that variations in RSN activation (and coactivation) may to a large part be driven by changes in the variance of the underlying BOLD signal. When using a short window and GSR, these values reduce to 0.43 on the group level and 0.51 (SE=0.30) on the single subject level. Again, this reduction is due to GSR and not due to the shorter window (see table S3).

### 3.3 During peaks in BOLD variance, the brain is closer to average FC structure

We asked the question what influence the modulations in instantaneous BOLD variance have on FC. We saw that these modulations are closely related to modulations in instantaneous average correlation. What is more, RSN activations are modulated in the same way. These windows are evident in the time courses as local maxima and we will refer to them as “peaks” in the remainder of the text. On the other hand, BOLD variance also possesses local minima where it drops below its mean; we will refer to these local minima as “troughs”.

Naively, we could think of three interpretations for these fluctuations:

1. During peaks in instantaneous BOLD variance, the strong fluctuations in the BOLD signal lead to a low signal to noise-ratio (SNR). In this case, the correlation structure would be largely destroyed and a noisy pattern of all-to-all-correlations would arise and explain the high values of instantaneous average correlation in these windows.
2. During peaks in instantaneous BOLD variance, SNR is high and the correlation structure is close to the average FC.
3. During peaks in instantaneous BOLD variance, SNR is high and dFC takes on highly structured patterns that deviate strongly from the average FC.

**Table 1:** Group level correlations between the three dFC measures for real data/surrogates with preserved long-term correlation/simulated data. For real data, correlation between dFC measures and RSN time courses (taking the sum at each point in time) is also given. All values are for w=120s and without GSR.

|  | inst.av.corr. | sim.to avFC | RSNs |
| --- | --- | --- | --- |
| inst.BOLD var. | 0.68/0.69/0.90 | 0.39/0.39/0.63 | 0.74 |
| inst.av.corr. |  | 0.36/0.31/0.59 | 0.67 |
| sim.to avFC |  |  | 0.32 |

As to hypothesis 3, we have already demonstrated that RSNs across time can be largely explained by the average FC. Thus this is not plausible. Hypothesis 1 does not explain the positive correlation of RSN time courses with instantaneous BOLD variance. The correlations mean that RSNs are strongly activated when instantaneous BOLD variance is high. To test hypotheses 1 and 2 against each other, we employ a third global measure of temporal dynamics: We measure, for each time window, the similarity of the dFCM to the avFCM by taking the correlation between the two. The correlations are summarized in table 1 and show that in line with hypothesis 2, there is a tendency for the dFCM to be more (rather than less, as hypothesis 1 stated above predicts) similar to the avFCM when instantaneous BOLD variance is high (correlation: 0.39), instantaneous average correlation is high (correlation: 0.36) and when RSN activity is high (correlation: 0.32). As before, we compare these group results to the single subject level and find that the mean correlation is about 0.40 for all pairs, with large standard errors of about the same size (see table S3 for exact numbers). This indicates again that there is large variability across subjects, however, the tendency is clear, and the alternative hypothesis that high instantaneous BOLD variance is equivalent to strong noise can be rejected. These results are reproduced for the shorter window and with GSR (SI results section S2.4 and table S3).

In order to further explore this, we plot in the left panel of figure 8 the joint distribution of instantaneous BOLD variance and similarity to avFC, averaged over all subjects. It is evident that this relationship is not linear. Furthermore, there seem to be very many time windows in which the instantaneous BOLD variance and the similarity to avFC are both very low (prominent cluster at the bottom left). In the next step, we will use surrogate data to test whether this observations really hints at nonstationarity, as could be suspected.

### 3.4 Stationary dynamics cannot fully reproduce distributions of modulations

It has been suggested that the modulations apparent in dFC are well within what is predicted by a multivariate Gaussian process, adjusted for the autocorrelations inherent in the BOLD signal (Handwerker et al., 2012; Hindriks et al., 2015), i.e. a stationary process as defined in the introduction. This is equivalent to the null hypothesis that observed modulations can be fully explained by the presence of long-term correlations and thus, by noisy fluctuations around the average FC structure. We test the hypothesis of stationarity by constructing appropriate surrogate data. We use a method that randomizes the phases, preserving the pairwise correlation structure and thus the the power spectrum (Handwerker et al., 2012; Hindriks et al., 2015). This results in any nonstationary properties being removed. Figure 9A and C shows an example - it is evident that also the surrogates possess the bands/events noted before. Thus, these synchronized movements are not per se an indication of any active process. Rather, they are the consequence of the average correlations/the 1/f power spectrum, which is shown in figure 9E to be reproduced by the surrogates.

**Figure 8:**
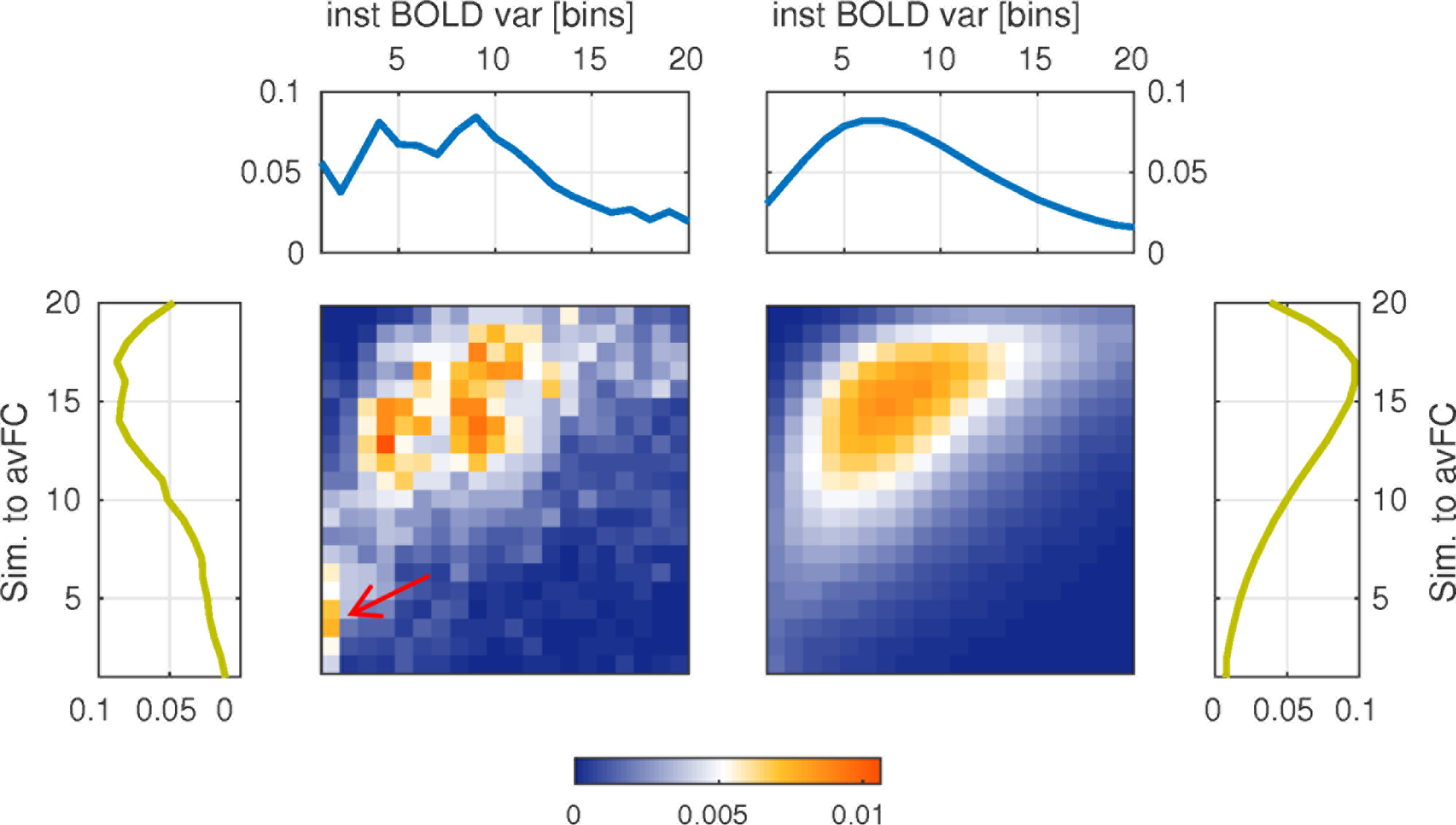
Joint distribution of instantaneous BOLD variance (x-axes) and similarity to avFC (y-axes) for empirical data (left) and surrogates (right). The marginal distributions are shown at the top and sides. The red arrow marks the significant (tested with surrogates) difference between the data and the surrogates (section 3.4).

First, we examine the correlations between the same three measures as before. Modulations are apparent also in the time courses derived from surrogate data, and the correlations between them are preserved (see table 1). This means that the prominent relationship between BOLD variance, average correlation and dFC structure is reproduced by stationary surrogate data.

Second, we examine the size of the peaks and troughs, i.e. local minima and maxima of the modulations. This has been the focus of many dFC studies and has been referred to in some instances as “hypersynchronization” (Hutchison et al., 2012). We note that these values differ greatly across subjects, meaning that, in absolute terms, their degree of variability is very heterogeneous. For example, the subject with the maximum range of instantaneous average correlation has a maximum of 0.78 and a minimum of 0.22; the one with the minimum range, a maximum of 0.40 and a minimum of 0.29. It has been suggested that intrinsic variability is meaningful (Garrett et al., 2010), but this topic is beyond the scope of this study. We conclude that under these circumstances, a group analysis averaging over the minimum/maximum values would not represent the subjects well. We instead report results for single subjects. We compute, for each subject, 100 sets of surrogate data. We find that they usually reach more extreme values than the real data, with the exception of only a few subjects. Thus, we do not find any signs of nonstationarity using this approach.

Third, in order to average across subjects, we introduce a normalization step which equalizes the “ranges” of each subject. We do this by computing histograms from the time courses and averaging the number of windows falling into each bin. Figure 10 shows the average distributions together with those obtained from the surrogates. Thereby, we ask whether the surrogate time courses are just scaled versions of the real data. Of course the minima and maxima are now the same for each subject. Therefore, we count the number of windows falling into the first and last bins of the distributions, therebey testing whether the time courses “visit” the peaks and troughs, respectively, more often than can be explained by the stationary surrogates. We find that the minimum bins are more populated in the real data than in the surrogates (none of the surrogate datasets has the same or a larger number of windows in the minimum bin). This means that real data possess more minima or remain there longer than the surrogates. This result holds for both window lengths and both with and without GSR (SI results section 2.5 and figures S22-24).

We note that also the shapes of the distributions for single subjects are very heterogeneous. Furthermore, for many subjects, the surrogate distributions do not seem to reflect the original data at all. We explain this by the fact that we are looking at very slow modulations and although our scanning time is quite long (22 minutes), we are possibly not sampling these distributions properly. Averaging over subjects seems to be the best choice in this situations. Analyzing multiple sessions obtained from the same subject might be an interesting extension of the analysis presented here.

Returning to figure 8 and now comparing the two panels (empirical vs. surrogates), it becomes clear that the modulations and the correlations between different measures are explained by the power spectrum and the presence of average correlations. At the same time, the number of windows that can be classified as troughs (arrow in figure 8) is *not* explained by these stationary features of the data.

**Figure 9:**
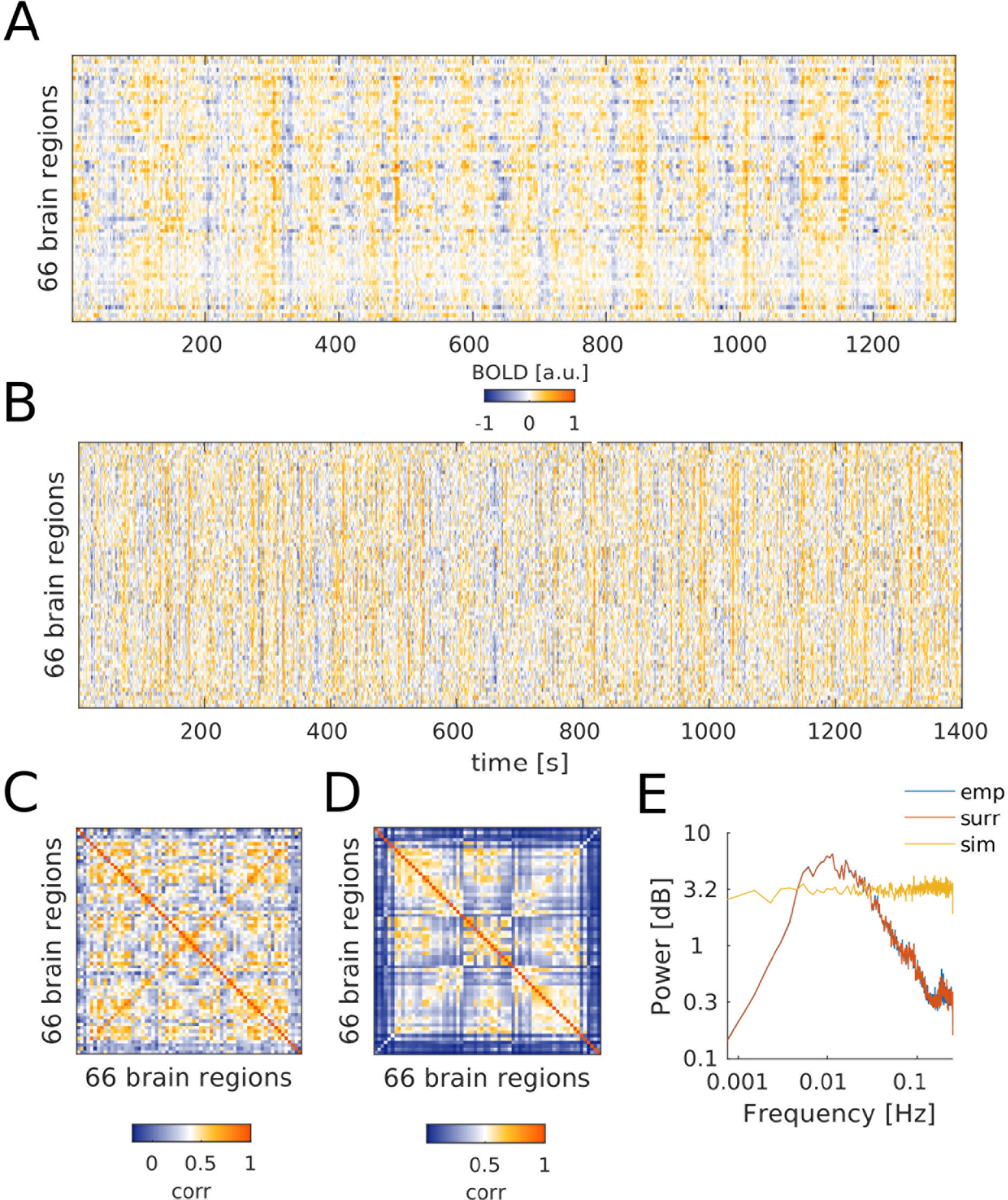
Stationary null data used in this study. **A** Phase randomized BOLD time courses from one subject. **B** Simulated data which were passed through the Balloon-Windkessel model. **C** Average FC matrix obtained from the surrogate time courses shown in **A**. It is nearly identical to that of the empirical data from which the surrogates where derived. **D** Average FC matrix obtained from the simulated time courses shown in **B**. The correlation between the two FC matrices is about 0.6. **E** Average power spectra of empirical, surrogate, and simulated data. Empirical and surrogate spectra lie almost exactly on top of each other.

**Figure 10:**
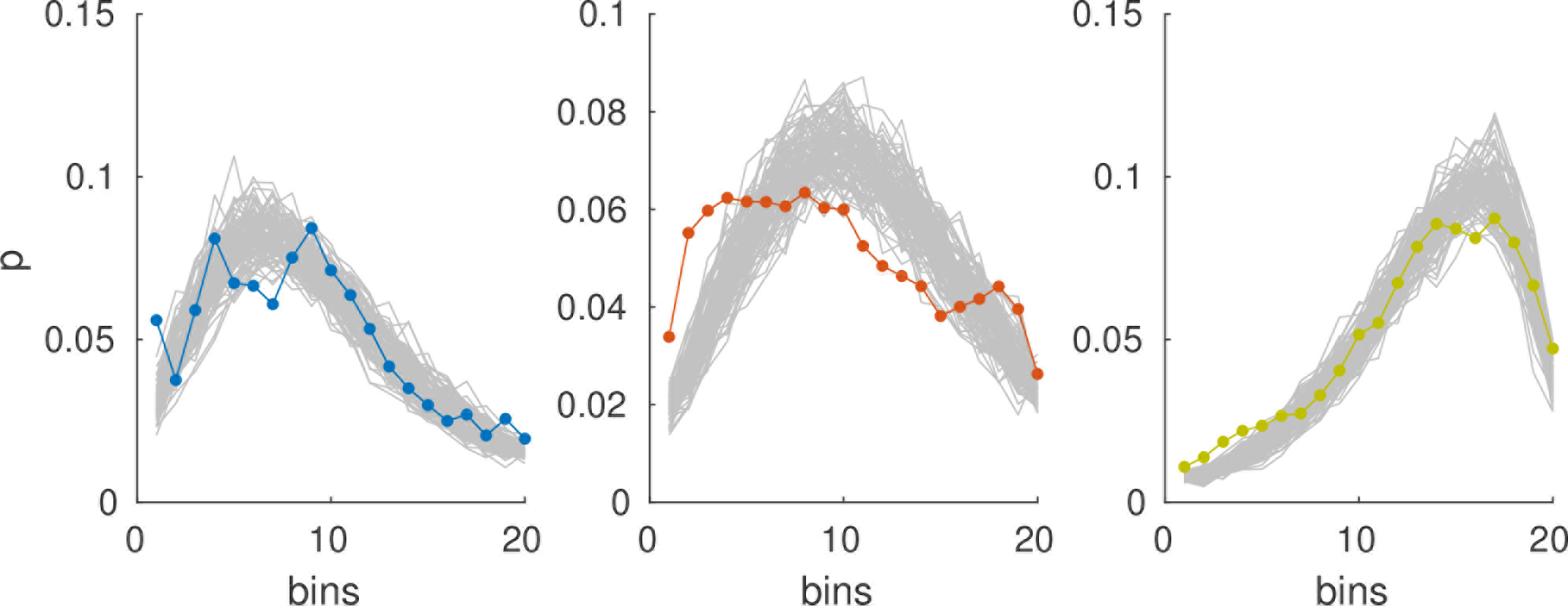
Comparison of empirical (colored lines) and surrogate (grey lines) histograms. Left panel: Instantaneous average correlation, middle panel: instantaneous BOLD variance, right panel: similarity to avFC.

### 3.5 RSN temporal dynamics in surrogates and stationary model

RSNs can be extracted in the exact same way from the surrogate as from the empirical data. When computing the histograms of the contributions in order to evaluate the propensity for coactivation (figure 6B), we find again strong temporal overlap as in the empirical data. In concordance with what was found in the previous section, the peak to the left is missing, i.e. what is reduced in the surrogate RSN time courses are periods in which networks are inactive. The same results are shown for the shorter window and when using GSR in the middle panels of figures S16-S18.

Our next step is to explicitly test the hypothesis that the observed modulations can be explained by correlated fluctuations around a single single fixed point - that is, by the average functional connectivity. We use a dynamic mean field model which possesses exactly this structure and which has been shown before (Glomb et al., 2017) to reproduce RSNs. Nodes in the model, representing brain regions, are connected according to effective connectivity obtained by combining diffusion tensor imaging results with a simple model (Gilson et al., 2016), see section 2.6.

Figure 9B and D show an example for a simulated signal and the average FC matrix computed from it. Figure 9E also shows the average power spectrum of all simulations. Clearly, while the correlation structure is captured well, the power spectrum is consistent with white noise, while empirical (and thus surrogate) data exhibit a 1/f spectrum (Biswal et al., 1995; Niazy et al., 2011).

As for the empirical data and the surrogate data, we compute time courses of our three global measures of temporal dynamics, and find that they exhibit very similar properties in terms of correlations between them (table 1). Likewise, extracting RSNs from the simulated data and computing the coactivation histograms (figure 6C) shows again strong coactivation, but fewer windows in the minimum bins of the distributions, even more prominently than in the surrogate data. It is important to note that the simulated data do not possess the same kind of variability as the empirical data as the simulations are done with the average connectivity matrix and merely reproduces the quantity of data by repeating the same (randomly initialized) simulation *S* times. This explains the differences apparent between surrogates, which are a true representation of the subject pool, and the simulations.

It is interesting to note that the 1/f power spectrum is in fact not necessary to explain these properties.

In conclusion, we have shown that

1. the modulations observed in BOLD temporal dynamics and RSNs are *stereotypical* in the sense that peaks correspond to strong RSN coactivation, high BOLD variance, high average correlation and high similarity of dFC to avFC (as opposed to the notion that each peak represents a different state),
2. the presence and the size of these modulations are explained by the average correlations present in the data, as shown by using surrogates and a stationary dynamic mean field model,
3. nonetheless minima of the modulations are visited more frequently than can be explained by stationary dynamics, suggesting the presence of an active, nonstationary process.

## 4 Discussion

This study does not explore the question whether the brain is nonstationary - it would be hard to find anyone who would not answer this question with “yes”. Instead, we are investigating whether “states” in the form of global dFC patterns obtained from fMRI BOLD signals can capture this nonstationarity. By investigating variances, correlation strenghts and patterns, we probe the second order statistics of the data, which are expected to be constant if the generative process is stationary. We are constrained by the data and methods available to us - limits on the number of subjects, scanning duration, as well as number of sessions; the temporal resolution which at a TR of 2s allows us to consider only frequencies below 0.25 Hz; the spatial resolution, though affording relatively precise source localization (for a global, noninvasive method), is not fully used because computational (for simulations and tensor construction/-decomposition) and statistical power are limited as well (this is why we only find 4-6 RSNs). Therefore, all the results presented specifically speak to the methods employed.

We used three simple measures to characterize modulations in the BOLD signal and their relationship to dFC: (1) instantaneous BOLD variance, (2) instantaneous average correlation, (3) similarity to avFC. We related these measures to RSN activation time courses obtained from empirical data via tensor decomposition. We found that a good part of the modulations we observe can be described as stereotypical events which evolve from a state where all three measures and RSN activations are low, to a state where these measures are high. Additionally to these descriptive results, we used surrogate data from which nonstationarities were removed, as well as simulated data obtained from a stationary model, to investigate the mechanisms behind our observations. We showed that many statistical properties of spontaneous fluctuations in the resting state BOLD signal are explained by the presence of long-term correlations and the characteristic power spectrum of these data. However, the dynamics of the transitions between peaks and troughs of the modulations found in our measures and in RSN time courses exhibit nonstationary features on the group level: There are more windows classified as troughs than can be explained by the stationary surrogates and simulated data. We find that there is a large variability across subjects, and explaining and modelling this variability is an interesting topic for future investigations.

### 4.1 Relationship between changes in activity and in FC: sliding windows vs point process

We have shown very clearly that there is a strong positive correlation between changes in BOLD variance and changes in FC (section 3.2), in parallel to findings on a finer spatial and temporal scale (Chawla et al., 1999; De La Rocha et al., 2007) and for phase relationships (Daffertshofer and van Wijk, 2011). While it may be clear that this relationship is expected in the sense that it is explained by the long-term correlations, it is not necessarily intuitive or without alternative. From recent work (Tagliazucchi et al., 2012; Griffa et al., 2017), it has become clear that dFC can be studied converting the BOLD signal into a point process, binarizing using the time points at which the signal crosses a threshold given by its standard deviation is crossed. FC in this context is defined as simultaneous crossing of the threshold, and surprisingly few time points are necessary to recover RSNs and rich spatiotemporal patterns. This relates to our result shown in section 3.3 that stronger BOLD variance occurs together with stronger expression of FC structure. In this framework, it is obvious that greater variance implies greater FC. This way, the present study explains why point process and sliding window-approaches to dFC have led to such similar results.

Considering these points, it should be clear that, when reporting changes in FC, for example due to task, disease, anaesthesia, etc., it should be carefully assessed which role the changes in activity play. In other words, changes in FC could just be “apparent” changes and not really indicate any difference in the communication between the areas in question. In general, while it is straightforward to define FC in terms of statistical dependency of two full time courses, it seems that to what exactly it refers in terms of dynamical changes is not as clear. Effective connectivity might be a useful complementary approach. To quantify the contribution of BOLD fluctuations to observed changes in FC, the surrogates used here could be useful because no real changes in FC occur. A suitable metric to quantify the distance between distributions of measures of dynamics, for example based on entropy, could capture the “residual” differences between surrogates and real data caused by real changes in FC.

### 4.2 Time scales and possible origins of fluctuations

One limitation of our methods here is that we cannot make clear predictions about the origin of the observed time scales: In the surrogate data, the Fourier spectra of the time courses are preserved as per construction, and the simulated data, while preserving many properties of the dynamics, do not exhibit the same kind of power spectrum at all (figure 9E). Recent work has begun to illuminate the relationship between network structure and power spectrum (He, 2014; Chaudhuri et al., 2016; Ocker et al., 2017). Elsewhere, it has been shown that slow (<0.1 Hz) fluctuations observed on the level of BOLD activity can arise from gamma band oscillations on the neuronal level (Cabral et al., 2011, 2014), and it has been suggested (Deco and Kringelbach, 2016) that they are a signature of long-range communication in the communication through coherence (CTC)-framework (Fries, 2005). Furthermore, animal (Shmuel and Leopold, 2008; Schölvinck et al., 2010; Thompson et al., 2013) and EEG/ECoG studies in humans (He and Raichle, 2009; Britz et al., 2010; Musso et al., 2010; Hiltunen et al., 2014) have shown a direct relationship between low (< 1 Hz) frequency electrical activity and dFC, sometimes manifestig in travelling waves (Thompson et al., 2014; Matsui et al., 2016). Therefore it seems that the observed fluctuations are at least partly of neural origin and evident in the sliding window time courses (Zalesky and Breakspear, 2015).

Fluctuations of our three global measures occur on an even slower time scale (≈0.01 Hz). The time scale reported here is consistent with previous findings (Hutchison et al., 2012; Hansen et al., 2014; Ponce-Alvarez et al., 2015), and in particular, modelling studies (Hansen et al., 2014; Ponce-Alvarez et al., 2015) have shown that they arise from spontaneous fluctuations in the band around 0.1 Hz and are slow modulations in the level of synchronization, manifesting as global state transitions or metastability. We used two different window lengths, showing that the properties in terms of second order statistics do not change much although windowing produces a new power spectrum that reflects modulations in power in the original data (Zalesky and Breakspear, 2015). For the shorter window, modulations in our dFC measures and RSN time courses have a shorter duration, suggesting that they are more narrowly localized in time than can be captured by slidng windows of either length. Using point processes (Tagliazucchi et al., 2012; Karahanoğlu and Van De Ville, 2015) is a promising alternative. At the same time, we investigated the impact of global signal regression, and although we could exclude the possibility that all our observations are due to modulations in the global signal strength, there were clear differences in terms of the clustering performance and RSN time courses. While our results further confirm that GSR is not beneficial for RS fMRI data analysis, it also remains unclear what the origin of this signal is.

### 4.3 RSNs overlap in time

Previous studies have suggested that RSN activation occurs as a sequence of transients (Baker et al., 2014; Ponce-Alvarez et al., 2015), putting the focus on their being separate in time. Here, we find that they coactivate a lot of the time, which was also reported by de Pasquale et al. (2012); Smith et al. (2012); Karahanoğlu and Van De Ville (2015). We find that RSNs are most strongly activated and coactivated during periods of high average correlation. This is at odds with the idea that each peak in the measures represents a distinct “state”, and suggests that, rather, we are looking at stereotypical events. This is in line with the interpretation given in Mitra et al. (2014): the activation of RSNs is embedded in stereotypical events triggered by global waves of BOLD activity. Matsui et al. (2016) found that in mice, typical co-activations between brain areas are embedded in global waves of activity, and that this is true both on the level of Calcium signals and the hemodynamic signal.

We note that we used different ways of computing pairwise FC for tensors/RSNs and for dFC measures: in the former case, we used mutual information, in the latter, Pearson correlation. MI has been shown to yield better clustering results, possibly because it is more robust against outliers (Glomb et al., 2017), and provides a naturally nonnegative input to the decomposition algorithm. Correlation, on the other hand, is the most common measure used in dFC studies. The fact that we nonetheless find close correlations between RSN time courses and correlation-based dFC measures speaks to the robustness of our results.

Our findings go hand in hand with the result that dFC structure is most of the time close to avFC and that most modulations can be explained by fluctuations around this average structure. While this may be surprising, it might indeed be a sign of optimal organization of the brain network, as suggested by recent work combining dFC with control theory (Gu et al., 2015; Betzel et al., 2016b). From the point of view proposed there, it is desirable to remain in a regime from which frequently visited states are easily reached, both in terms of energy cost and time. In this light, it would actully be surprising to see large excursions from the average during RS.

### 4.4 Conclusions and Outlook

The basic problem we are facing is that stationarity or nonstationarity are not properties of the *data*, but of the underlying *process* which generates the data. It is clear that we do not fully understand the mechanisms underlying the BOLD signal, let alone brain activity. Here, we have shown that whether we conclude that our data exhibits signs of nonstationarity depends on 1) our analysis methods: looking at the size of single events vs. looking at the distribution of these events, 2) whether we look at the group or the single subject level. Results seem to depend more strongly on the information available in the BOLD signal than details of our analysis as evident from the fact that results are robust against window length, GSR, and motion correction. On the group level, we find signs of nonstationarity - speculating, the high number of of windows in troughs could be seen as “echoes” of bistable dynamics in underlying neural activity. This result is not easily applied at the single subject level due to the great inter-individual variance. Combining BOLD data with other time scales, e.g. EEG, is very promising, and recent progress in source localization techniques might overcome the obvious problems with spatial resolution (Liu et al., 2017).

On the other hand, we have extracted some results that are interesting even though they are not directly linked to nonstationarity: 1) there is a tight relationship between BOLD variance and correlation, 2) RSNs coactivate strongly, suggesting that global dynamics of BOLD variance are a “background” to inter- and intra-network dynamics which cannot be “explained away” by the global signal, 3) specifically, correlation structure is more strongly expressed when BOLD variance is high. Taking the example of the DMF model used here, while it is stationary, it still possesses many interesting properties like two different time constants (excitatory and inhibitory pools) and an underlying effective connectivity structure which in itself is a complex network. What our results suggest is that the dFC community should be careful to define what “states” are when talking about “state switching” and implying nonstationarity in the sense of changing correlation structure over time because the richness of the dynamics within a stationary regime is not fully explored yet. Particularly, we still do not understand the relationship between structure and function although we do know that it is important. For example, the dynamic mean field model, which is a fairly complex model, while reproducing average FC quite well, does not reproduce the RSNs very well, and this cannot be attributed to the lack of nonstationarity as phase-randomized surrogate data do reproduce them.

We have to consider the possibility that we cannot observe the dynamics sufficiently well due to the limited data available per subject. Recent work (Finn et al., 2015) shows that FC can be used to identify single subjects. In this view, what we are analyzing here are only the patterns that are most common, and since the modulations are very slow, our sampling of the state space may not be sufficient (see sections 3.2 and 3.3).

The model used here is able to reproduce some of the hallmarks of dFC due to the scale-free temporal structure of BOLD signals (He, 2014), but lacks the non-stationarities as well as the interindividual differences. Possibly a model which has previously been linked to multistability at the level of neural fluctuations (Freyer et al., 2011), and which has recently been shown to reproduce features of fMRI RS temporal dynamics (Deco and Kringelbach, 2016), could be of interest. As before, they key is that the system is close to a bifurcation, in this case, a subcritical Hopf bifurcation. Cast in these terms, the system would be in a regime of damped oscillations when the global measures are low, and in an oscillatory regime when they are high; switching occurs due to noise. Another property that the DMF model used here does not reproduce is the power spectrum. We find that the time scale of modulations is not explained in spite of the fact that the hemodynamic signal is modelled explicitly (Friston et al., 2000), i.e. the 1/f power spectrum of real BOLD signals is not shared by the simulated data. Whether the Hopf model could reproduce the power spectrum should be included in the investigations. Furthermore, simulations using individual SC/EC should be helpful, and progress in fiber tracking techniques and model-based optimization hold promise in this point.

## Funding (authors’ initials given after grant numbers)

This work was supported by the European Union, FP7 Marie Curie ITN “INDIREA” (Grant N. 606901; KG), FP7 FET ICT Flagship Human Brain Project (Grant N. 604102; MG), ERC Advanced Human Brain Project (Grant N. 604102; GD), Horizon2020 ERC Consolidator (Grant N. …; PR);

the Spanish Ministry for Economy, Industry and Competitiveness (MINECO) project “PIRE-PICCS” (Grant N. PCIN-2015-079), SEMAINE ERA-Net NEURON Project (Grant N. PCIN2013-026; APA), and ICoBAM (Grant N. PSI2013-42091-P; GD);

the James S. McDonnell Foundation (Brain Network Recovery Group, Grant N. JSMF22002082; PR);

the German Ministry of Education and Research (Grant N. 01GQ1504A and 01GQ0971-5; PR);

the Max-Planck Society (Minerva Program; PR)

